# INTERACT: Interactome Network Targeting via Enzyme Reactivity and Activity-based Chemoproteomic Tools

**DOI:** 10.1101/2025.11.03.686322

**Authors:** Exequiel O. J. Porta, Zisis Koutsogiannis, Karunakaran Kalesh, Paul W. Denny, Patrick G. Steel

## Abstract

This chapter details an activity-based chemoproteomic methodology, INTERACT (Interactome Network Targeting via Enzyme Reactivity and Activity-based Chemoproteomic Tools), for the functional interrogation of host-pathogen interactions (interactome) in a physiologically relevant context. The exemplar protocol utilises a small, cell-permeable fluorophosphonate probe to covalently label active serine hydrolases simultaneously within *Leishmania mexicana* parasites and their murine macrophage host cells during active infection. Subsequent biorthogonal click chemistry, affinity enrichment, and quantitative mass spectrometry using Tandem Mass Tags (TMT) enable the identification and relative quantification of functionally active enzymes across the interactome. This method facilitates the delineation of dynamic changes in enzyme activity during the infection process, yielding rich insight into host–pathogen biochemical networks, including the identification of pathogen virulence factors and host responses. Therefore, INTERACT delivers a robust and quantitative workflow to probe enzymatic activity across two species in a single experiment under native infection conditions, without the need for genetic manipulation, providing an invaluable platform for dissecting pathogenesis and uncovering novel therapeutic targets in infectious diseases.

## 1. Introduction

The molecular crosstalk between pathogens and their hosts is a dynamic and complex process that is fundamental to the progression of infectious diseases. Understanding this intricate process is critical for deciphering mechanisms of pathogenesis and identifying novel therapeutic targets (Dix et al., 2016). The intracellular parasite *Leishmania*, the causative agent of leishmaniasis, provides a compelling model system (Isern et al., 2025). This parasite has a digenetic lifecycle, alternating between a motile promastigote form in the insect vector and a non-motile amastigote form that resides and replicates within host macrophages. The survival and proliferation of the parasite within the hostile environment of the macrophage’s parasitophorous vacuole depend entirely on its ability to adapt to and manipulate the host cell through a complex interplay of parasite virulence mechanisms and host cell responses (Liu and Uzonna, 2012; Gupta et al., 2013; Costa-da-Silva et al., 2022).

Traditional methods for studying interactomes (such as yeast two-hybrid and co-immunoprecipitation) and conventional proteomics, have significant limitations in this context (Ito et al., 2001; Maccarrone et al., 2017; Wu et al., 2025). They often require protein overexpression or cell lysis, both of which disrupt the native host–parasite context and cellular architecture, thereby perturbing the interactome. They also tend to miss transient or weak interactions and report protein abundance rather than functional activity, yielding an incomplete picture of the dynamic cellular processes at play during infection.

Activity-based protein profiling (ABPP) has emerged as a powerful chemical proteomics strategy that overcomes these challenges (Fang et al., 2021; Porta, 2023). ABPP utilizes small-molecule chemical probes to directly measure the functional state of entire enzyme families within complex biological systems, including living cells (Cravatt et al., 2008; Porta and Steel, 2023). Crucially, this approach provides an *in situ* snapshot of enzymatic activity and, because labelling can be performed in live or otherwise intact (non-lysed) cells with cell-permeable probes, preserves the cellular context of host–pathogen interactions (Kovalyova and Hatzios, 2019; Kozoriz and Lee, 2025). By employing broad-spectrum activity-based probes (ABPs) that target entire enzyme families, ABPP enables exploration of the interactome at the network level.

This chapter describes INTERACT (**I**nteractome **N**etwork **T**argeting via **E**nzyme **R**eactivity and **A**ctivity-based **C**hemoproteomic **T**ools), an ABPP workflow designed to map changes in enzyme activity that occur upon pathogen invasion of a host system. Whilst this chapter presents INTERACT in the context of the role of serine hydrolases in the *Leishmania*–host interactome during infection, the workflow is, with system-specific optimisation, adaptable to other enzyme activities and host–pathogen models. The key reagent is an activity based-probe (ABP), i.e., a chemical probe, that is capable of crossing multiple membranes and labels enzymes in both the intracellular pathogen and host cell simultaneously. In this specific example, the ABP is a small, cell-permeable fluorophosphonate (FP) probe that labels the serinome (i.e., the set of serine hydrolase enzymes/activities detected by ABPs) in both intracellular parasites and host macrophages *in situ* (Isern et al., 2025). Here, “serinome” denotes the set of serine-hydrolase enzymes/activities detected by FP-ABPP.This live-cell-first strategy offers a key advantage over lysate-only approaches that often depend on larger, less permeable probes (Porta et al., 2022). After covalent labelling and cell lysis, a biorthogonal Copper (I)-Catalysed Azide–Alkyne Cycloaddition (CuAAC) reaction is used to append a biotin tag to the probe’s alkyne handle, enabling stringent affinity enrichment prior to MS analysis. In practice, infected macrophages are treated with the alkyne-bearing ABP, which covalently captures catalytically competent serine hydrolases across both proteomes, minimising artefacts from cell disruption and preserving native context. Following enrichment, probe-labelled proteins are identified and quantified by LC-MS/MS. A defining feature of INTERACT is the use of Tandem Mass Tag (TMT) multiplexing, which enables parallel comparison of multiple conditions (e.g., ±probe, ±inhibitor) and relative quantification of enzyme activities. Inclusion of no-probe channels helps discriminate true targets from background. The protocol that follows (**Figure 1**) provides step-by-step guidance from culture and infection to data analysis for functional interrogation of the host–pathogen interface in this context.

**Figure 1.**
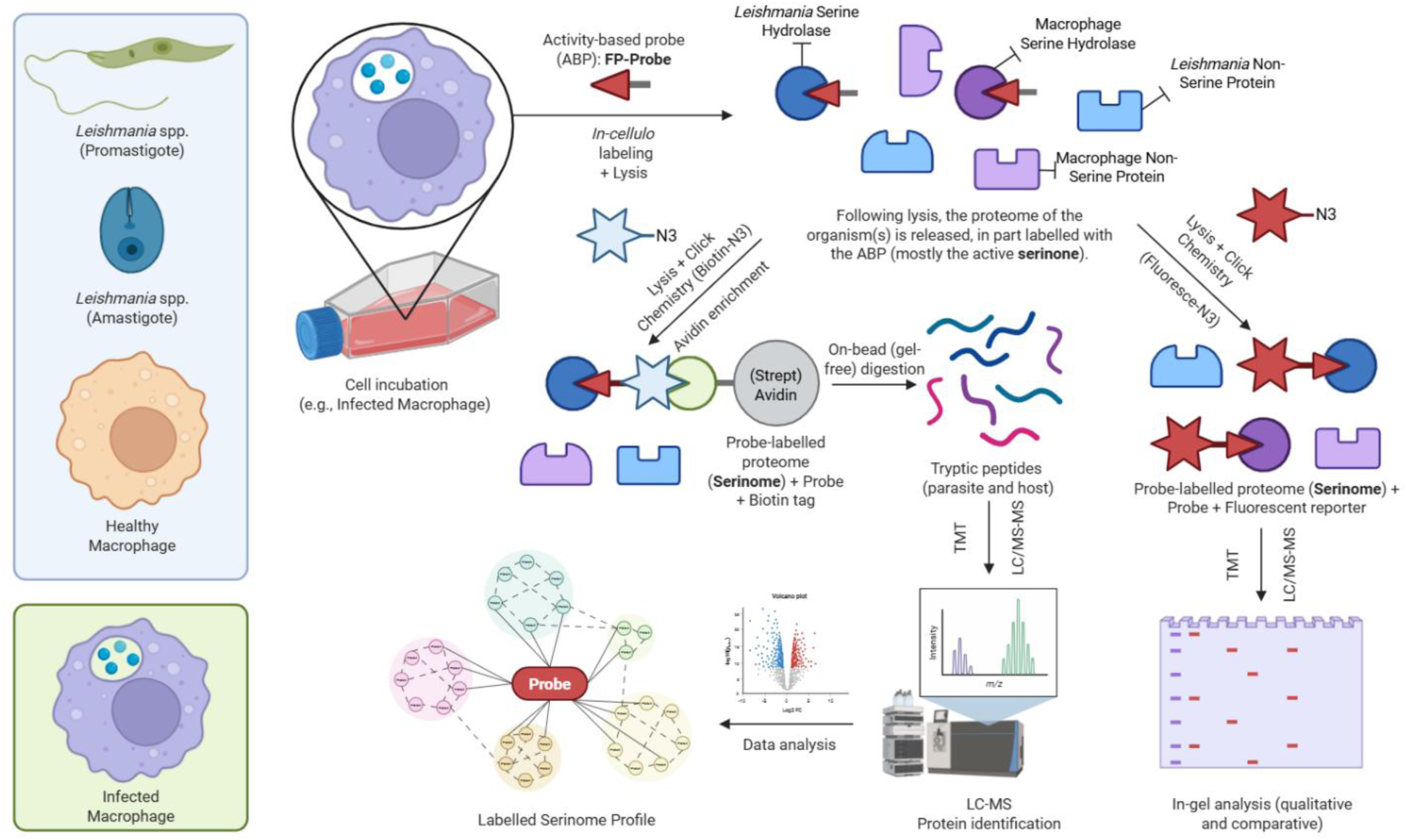
INTERACT ABPP/TMT workflow for (mono- and) dual-proteome profiling. Each sample type is labelled in separate incubations: (i) *Leishmania* metacyclic promastigotes, (ii) axenic amastigotes, (iii) uninfected RAW 264.7 macrophages (**Left**, blue box), and (iv) infected RAW 264.7 macrophages (**Left**, green box). After independent lysis (**Top**), probe-labelled proteins (in this case the serinome) undergo CuAAC to azide reporters (fluorophore or biotin) for either in-gel visualisation (SDS–PAGE, **Right**) or streptavidin-based enrichment (**Centre**), followed by on-bead digestion. Peptides from each sample are then TMT-labelled and pooled only at this step for LC–MS/MS, enabling identification and relative quantification of active enzymes and their regulation across conditions (**Bottom**).

## 2. Materials

Use high-purity reagents and maintain sterile technique for all cell culture steps. Prepare all buffers and stock solutions in advance (*see* Section “Stock Solutions”) and keep them on ice or at appropriate storage conditions until use. Prepare all solutions using either Milli-Q water (for biological solutions) or LC–MS-grade water (Thermo Fisher Scientific, catalogue number 51140) and analytical grade reagents, unless otherwise specified. Prepare and store all reagents according to the instructions below. Follow all institutional waste disposal regulations.

### 2.1 Reagents

**Table 1** outlines the key materials required for the INTERACT protocol, organised by reagent type, consumables, equipment, and stock solutions. Supplier catalogue numbers are provided where available.

**Table 1.** Key materials required for the INTERACT protocol.

| Reagent | Supplier | Catalogue Number |
| --- | --- | --- |
| *Leishmania mexicana*<br>promastigotes | In House | MNYC/BZ/62/M379 |
| RAW 264.7 (TIB-71, ATCC)<br>murine macrophage cells | In House | TIB-71 |
| Schneider's Drosophila Medium | Thermo Fisher Scientific | 21720024 |
| Fetal Bovine Serum (FBS),<br>Qualified | Thermo Fisher Scientific | 16000044 |
| Penicillin-Streptomycin Solution<br>(100X) | Thermo Fisher Scientific | 15140122 |
| Dulbecco's Modified Eagle's<br>Medium (DMEM) | Thermo Fisher Scientific | 11965092 |
| Dulbecco's Phosphate-Buffered<br>Saline (PBS), pH 7.4 | Thermo Fisher Scientific | 14190144 |
| Ethyl [5-(prop-2-yn-1-yloxy)pentyl]<br>phosphonofluoridate (**FP Probe**)<br>(**Figure 2**) | **See Note 1** | **See Note 1** |
| Dimethyl sulfoxide (DMSO),<br>Anhydrous | Sigma-Aldrich | 276855 |
| Complete Mini, EDTA-free<br>Protease Inhibitor Cocktail | Roche | 1183170001 |
| Pierce Rapid Gold BCA Protein<br>Assay Kit | Thermo Fisher Scientific | A53226 |
| Biotin-Azide (PEG<sub>4</sub> Azido-Biotin) | Thermo Fisher Scientific | B10184 |
| Copper(II) Sulfate, pentahydrate<br>(CuSO<sub>4</sub>·5H<sub>2</sub>O) | Sigma-Aldrich | C7631 |
| Tris(benzyltriazolylmethyl)amine<br>(TBTA) | Sigma-Aldrich | 678937 |
| Sodium Ascorbate | Sigma-Aldrich | A4034 |
| Methanol (MeOH), LC-MS Grade | Thermo Fisher Scientific | A456-4 |
| Sodium Dodecyl Sulfate (SDS) | Sigma-Aldrich | L3771 |
| NeutrAvidin-Agarose Resin | Thermo Fisher Scientific | 29200 |
| Urea | Sigma-Aldrich | U5378 |
| Triethylammonium Bicarbonate (TEAB) Buffer, 1 M, pH 8.5 | Thermo Fisher Scientific | 90114 |
| Tris(2-carboxyethyl)phosphine (TCEP) Hydrochloride | Thermo Fisher Scientific | 20490 |
| Iodoacetamide (IAA) | Sigma-Aldrich | I1149 |
| Trypsin Protease, MS Grade | Thermo Fisher Scientific | 90057 |
| TMT10plex Isobaric Label Reagent Set | Thermo Fisher Scientific | 90110 |
| Hydroxylamine, 50% Solution | Thermo Fisher Scientific | 90115 |
| Acetonitrile (ACN), LC-MS Grade | Thermo Fisher Scientific | 51101 |
| Formic Acid (FA), LC-MS Grade | Thermo Fisher Scientific | 85178 |
| Pierce Peptide Desalting Spin Columns, 10-30 μL | Thermo Fisher Scientific | 89851 |

### 2.2 Consumables

1. T-25 and T-75 culture-treated flasks with vented caps.
2. 1.5 mL and 2.0 mL low-protein-binding microcentrifuge tubes.
3. Filtered pipette tips (10 µL, 200 µL, 1000 µL). Use low-bind tips for peptide handling post-digestion.
4. Cell scrapers.
5. Serological pipettes (5 mL, 10 mL, 25 mL).

### 2.3 Equipment

1. Humidified incubator with 5% CO_2_, set to 37°C.
2. Incubator set to 26°C.
3. Class II biosafety cabinet.
4. Inverted microscope for cell culture monitoring.
5. Refrigerated microcentrifuge capable of ≥16,000×g.
6. Benchtop centrifuge with swinging-bucket rotor for cell culture plates/tubes.
7. End-over-end rotator.
8. Thermomixer or heating block capable of maintaining 30°C, 37°C, and 95°C.
9. Vortex mixer.
10. Vacuum concentrator (e.g., SpeedVac). Should be capable of drying samples without heating above ∼30°C (to avoid peptide degradation).
11. Spectrophotometer or microplate reader capable of measuring absorbance at 562 nm for BCA assay.
12. Orbitrap Ascent Mass Spectrometer (Thermo Fisher Scientific) or equivalent high-resolution Orbitrap instrument.
13. Thermo Scientific Ultimate 3000 RSLCnano UHPLC system or equivalent nano-flow liquid chromatography system.
14. Computer with data analysis software.

### 2.4 Stock Solutions, Buffers and Growth Media

1. Cell culture Growth Medium: DMEM (with 4.5 g/L glucose, L-glutamine and pyruvate) supplemented with 10% (v/v) FBS and 1% (v/v) Penicillin-Streptomycin. Prepare fresh and store at 4°C for up to 4 weeks. Warm to 37°C before use.
2. Promastigote Growth Medium: Schneider’s Drosophila medium supplemented with 15% (v/v) FBS and 1% (v/v) Penicillin-Streptomycin. Store at 4°C.
3. Metacyclic Generation Medium: Schneider’s Insect medium supplemented with 20% (v/v) FBS, pH adjusted to 5.5. Store at 4°C. Briefly, to 500 mL Schneider’s Drosophila Medium, add 100 mL heat-inactivated FBS (final 20%) and 5 mL penicillin– streptomycin (100× stock). Adjust pH to 5.5 with 1 N HCl. Sterile-filter if pH was adjusted or if medium was prepared from powder.
4. Lysis Buffer: 25 mM Tris-HCl (pH 7.4), 150 mM NaCl, 1% (v/v) Triton X-100, 5% (v/v) glycerol. Prepare 50 mL in Milli-Q water. Store at 4°C for up to 1 month. Immediately before use, add one tablet of Complete Mini, EDTA-free Protease Inhibitor Cocktail per 10 mL of buffer.
5. FP Probe (**Figure 2**) Stock Solution (10 mM): Prepare in anhydrous DMSO. Aliquot into single-use volumes to avoid freeze-thaw cycles and store at -20°C, protected from light, for up to 6 months.
6. Biotin-N_3_ Stock Solution (5 mM): Prepare in anhydrous DMSO. Aliquot and store at -20°C.
7. Click Chemistry Stock Solutions: Prepare fresh for each experiment.
8. 100 mM CuSO_4_: Dissolve 25 mg of CuSO_4_.5H_2_O in 1 mL of LC-MS grade water.
9. 100 mM Sodium Ascorbate: Dissolve 19.8 mg in 1 mL of LC-MS grade water.
10. 10 mM TBTA: Dissolve 8.6 mg in 1 mL of DMSO.
11. Wash Buffer 1 (0.5% SDS/PBS): Add 5 mL of 10% (w/v) SDS solution to 95 mL of 1X PBS.
12. Wash Buffer 2 (6 M Urea/PBS): Dissolve 36.04 g of urea in 1X PBS to a final volume of 100 mL. Gentle warming may be required.
13. 100 mM TEAB Buffer: Dilute 1 M TEAB stock 1:10 with LC-MS grade water.
14. 50 mM TEAB Buffer: Dilute 100 mM TEAB 1:1 with LC-MS grade water.
15. 10 mM TCEP Solution: Prepare fresh. Dissolve 1.43 mg of TCEP-HCl in 500 µL of 50 mM TEAB.
16. 15 mM Iodoacetamide (IAA) Solution: Prepare fresh. Dissolve 2.78 mg of IAA in 1 mL of 50 mM TEAB. Protect from light.
17. 5% Hydroxylamine Solution: Prepare fresh. Dilute 50% stock solution 1:10 with 100 mM TEAB.
18. Trypsin Solution (0.2 µg/µL): Resuspend 4 µg of sequencing-grade trypsin in 20 µL of 100 mM TEAB immediately before use.
19. LC Solvent A: 0.1% (v/v) Formic Acid in LC-MS grade water.
20. LC Solvent B: 0.1% (v/v) Formic Acid in Acetonitrile.

## 3. Methods

The following sections describe a step-by-step protocol for applying the INTERACT method to profile active serine hydrolases in a *Leishmania*-infected macrophage system. Perform all cell culture steps in a sterile environment (Class II biosafety cabinet) and wear appropriate personal protective equipment, as *Leishmania* is pathogenic. For clarity, the procedure is divided into logical subsections. Steps within each subsection are numbered. Notes corresponding to specific steps are provided in the *Notes* section. Perform each experiment in at least biological duplicate, ideally triplicate.

**Figure 2.**
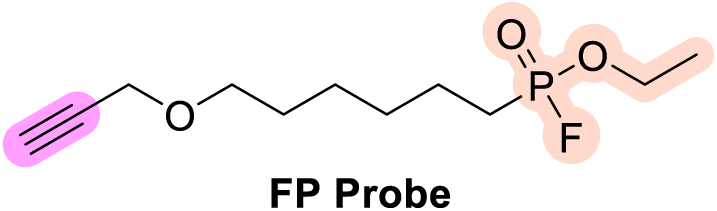
Structure of the cell-permeable alkyl-fluorophosphonate probe, ethyl [5-(prop-2-yn-1-yloxy)pentyl] phosphonofluoridate (**FP-Probe**). The FP warhead covalently reacts with active-site serines in serine hydrolases; the short alkyl linker terminates in an alkyne handle for CuAAC tagging after labelling. Colour legend: terminal alkyne handle in magenta; FP warhead in peach; alkyl linker and ether in black. This small, membrane-permeable probe (used at 10 µM in this protocol) supports *in situ* dual-proteome profiling of host macrophages and intracellular *Leishmania* during infection.

### 3.1 Culture of RAW 264.7 Macrophages and *Leishmania mexicana* Promastigotes

1. Culture RAW 264.7 macrophages in 10-12 mL of Growth Medium in T-75 flasks at 37°C in a humidified incubator with 5% CO_2_. Subculture the cells every 2–3 days to avoid stress and ensure optimal viability. Cell viability can be assessed using the Trypan Blue Exclusion Assay. Passage cells as needed to obtain sufficient numbers for the appropriate multiplicity of infection (MOI) and experimental depth.
2. In a 25 cm^2^ vented flask, inoculate *L. mexicana* promastigotes (strain MNYC/BZ/62/M379) into 5–10 mL Schneider’s medium (pH 7.0) with 15% FBS and 1% pen/strep. Incubate at 26°C (no CO_2_ needed). Start with an inoculum of ∼1 × 10^6^parasites/mL and allow them to reach late-log phase (∼1–5 × 10^7^/mL, typically 3–4 days).
3. Maintain *L. mexicana* promastigotes at 26°C in Promastigote Growth Medium in T-25 flasks.
4. To generate infective-stage metacyclic promastigotes, transfer late log-phase parasites into Metacyclic Generation Medium at a density of 5×10^5^ parasites/mL and incubate for 6-7 days at 26°C (*see* **Note 2**).

### 3.2 Macrophage Infection with *Leishmania*

1. Harvest RAW 264.7 cells from culture flasks using a cell scraper with gentle, careful motions to minimise physical damage. Count cells and assess cell viability via Trypan Blue exclusion assay, confirming a viability of at least 95%. Seed macrophages into an appropriate vessel and leave overnight to adhere at 37°C, 5% CO_2_.
2. On the day of infection, count the metacyclic promastigotes using a haemocytometer or cell counter. Aim to have an excess of parasites available to achieve the desired MOI.
3. Prepare control samples: Collect aliquots of non-infected macrophages and of the *Leishmania* promastigote culture to serve as experimental controls. Maintain these under the same conditions as the infected samples to allow direct comparison during downstream analyses (*see* **Note 3**).
4. Immediately before infection, confirm that monolayers are ∼80% confluent, healthy, and well-adherent under the microscope.
5. Inoculate with *Leishmania* (MOI 10:1): Calculate the volume of promastigote culture needed to achieve a MOI of 10 parasites per 1 macrophage (*see* **Note 4**). For example, for 5 × 10^5^ macrophages, use 5 × 10^6^ promastigotes. Replace macrophage media with fresh warm DMEM (enough to cover cells), then add the promastigotes directly to the well (or flask). Gently swirl to evenly distribute parasites.
6. Incubate for infection: Return the co-culture to 37°C, 5% CO_2_ for 4 hours. During this time, parasites will attach to and be phagocytosed by macrophages (*see* **Note 5**).
7. Wash to remove extracellular parasites: After 4 h, carefully aspirate the media, which contains non-internalised promastigotes. Gently wash the macrophage monolayer 3-4 times with appropriate volume of pre-warmed (37°C) 1X PBS to remove remaining extracellular parasites. For T-75, 3 × 5–10 mL PBS (37°C); inspect microscopically to ensure minimal free promastigotes. Washing steps are critical to ensure that subsequent analyses focus on macrophage-associated (internalised) parasites and not free promastigotes (*see* **Note 6**).
8. Add pre-warmed DMEM and immediately transfer the infected cells to a 37°C incubator for ∼15 min to adjust the temperature prior to probe labelling. In general, perform labelling at the optimal temperature for the system.

### 3.3 Chemical Probe Labelling of the Interactome

1. Prepare a working solution of **FP Probe** by diluting the 10 mM stock solution in pre-warmed (37°C) culture medium to a final concentration of 10 µM. In parallel, prepare a vehicle control by adding the same final concentration of DMSO without probe to the medium (*see* **Note 7**).
2. Add the probe solution (or the DMSO, no probe, control) directly to the infected cell culture: Remove the equilibration medium from the infected macrophages (from Step 3.2.8) and gently overlay the cell monolayer with the ABP-containing medium. Ensure full coverage of the cells. Minimise disturbance to avoid dislodging infected macrophages.
3. Incubate the cells with the probe for 1 hour at 37°C (*see* **Note 8**). During this period, the cell-permeable FP probes will penetrate both macrophages and parasites, covalently reacting with active serine hydrolases in each. Shield the samples from light if the probe is light-sensitive (*see* **Note 9**).

### 3.4 Cell Harvesting and Lysis

1. After 1 h probe incubation, immediately place the culture dish (or flask) on ice for handling and, gently, aspirate the medium (contains probe, so treat as chemical waste). Add 12 mL of ice-cold PBS carefully without disturbing the monolayer to wash the cells (to remove residual media and probe) and aspirate again. Repeat this step three times.
2. Add 10 mL of cold 1X PBS and use a cell scraper to detach the cells (*see* **Note 10**). Transfer the cell suspension to a pre-chilled 15 mL conical tube.
3. Pellet the cells by centrifugation at 500×g for 5 minutes at 4°C.
4. Carefully aspirate and discard the supernatant. Resuspend the cell pellet in an appropriate volume (e.g., 200–500 µL) of ice-cold Lysis Buffer freshly supplemented with protease inhibitors (*see* **Note 11**). Pipette up and down to resuspend the pellet. Transfer the cell suspension to a pre-chilled 1.5 mL microcentrifuge tube.
5. Agitate the lysate for 30 minutes at 4°C on an end-over-end rotator to ensure complete lysis. This allows efficient solubilization of proteins from both host and parasite while preserving enzyme–probe adducts.
6. Clarify the lysate by centrifugation at 16,000×g for 20 minutes at 4°C.
7. Carefully transfer the clarified supernatant (total protein lysate) to a new pre-chilled, low-bind microcentrifuge tube on ice, taking care not to disturb the insoluble pellet. If any particulate carryover is observed, re-centrifuge and transfer the clear supernatant to a fresh tube. This supernatant contains the probe-labelled proteins (the “interactome” proteome) (*see* **Note 12**). **STOP POINT**.

### 3.5 Protein Quantification

1. Determine the protein concentration of the clarified lysate using the Pierce Rapid Gold BCA Protein Assay Kit, following the manufacturer’s protocol.
2. Normalise the protein concentration of all samples to 1.5 mg/mL using Lysis Buffer.

STOP POINT.

### 3.6 Biorthogonal Cu-Catalysed Cycloaddition

1. In a chemical fume hood, set up the CuAAC reaction in each protein sample. For each 100 µL of normalised protein lysate (at 1.5 mg/mL), add the following reagents (final concentrations in parentheses): biotin–PEG_4_–azide 1 µL of 5 mM stock (50 µM), CuSO_4_ 1 µL of 100 mM stock (1 mM), TBTA ligand 1 µL of 10 mM stock (0.1 mM), and sodium ascorbate 1 µL of 100 mM stock (1 mM). Mix the components in order (copper last or ascorbate last according to preference; here adding ascorbate last initiates the reaction) (*see* **Note 13**). Sample volume can be less than 100 µL (*see* **Note 14**)
2. Incubate the reaction mixture (in dark) at room temperature for 1 hour, with periodic gentle mixing (*see* **Note 15**).

### 3.7 Protein Precipitation and Affinity Enrichment

1. Precipitate total protein by adding 9 volumes of ice-cold methanol to the lysate post-click reaction. Vortex and incubate at –80°C for ≥ 1 hour (or overnight) to precipitate proteins and remove excess unreacted reagents (*see* **Note 16**).
2. Centrifuge at 10,000×g for 10 minutes at 4°C to pellet the protein.
3. Wash the pellet twice with 1 mL of ice-cold methanol. Air-dry the pellet for 30 minutes at room temperature (*see* **Note 17**). **STOP POINT**.
4. Redissolve the protein pellet in a minimal volume of 2% SDS in PBS, just enough to dissolve it (e.g. 30–50 µL), then immediately dilute with PBS to a final SDS concentration of 0.1% (*see* **Note 18**).
5. While proteins are dissolving, equilibrate NeutrAvidin agarose beads. Gently resuspend the bead slurry (which may have settled) and transfer ∼50 µL of bead slurry per sample into a separate tube. Wash the beads 3 times with 4 volumes of 0.1% SDS in PBS (wash: add buffer, invert to mix, pellet beads by gentle centrifugation ∼1,000 × g 2 min, remove supernatant). After the final wash, remove supernatant.
6. Add 50 µL of washed NeutrAvidin-Agarose bead slurry to each sample and incubate for 2 hours at room temperature with end-over-end rotation (*see* **Note 19**).
7. Pellet the beads by centrifugation (1,500×g for 2 min) and discard the supernatant.
8. Remove supernatant (which contains non-biotinylated proteins – these can be discarded). Wash the beads with the following series to reduce non-specific binders (*see* **Note 20**):
9. 3 × 500 µL of 0.5% SDS in PBS (*see* **Note 21**).
10. 3 × 500 µL of 6 M urea in PBS (*see* **Note 22**).
11. 3 × 500 µL of PBS (*see* **Note 23**).
12. 1 × 500 µL of 50 mM TEAB (*see* **Note 24**).
13. After the final TEAB wash, remove as much liquid as possible.

### 3.8 On-Bead Reduction, Alkylation, and Tryptic Digestion

1. Resuspend the washed beads in 200 µL of freshly prepared 10 mM TCEP in 50 mM TEAB. Incubate for 45 minutes at 30°C to reduce disulfide bonds.
2. Wash the beads once with 400 µL of 50 mM TEAB. Centrifuge at 1,500×g for 2 min and discard the supernatant.
3. Resuspend the beads in 200 µL of freshly prepared 15 mM iodoacetamide in 50 mM TEAB. Incubate for 45 minutes at room temperature in the dark to alkylate free cysteines.
4. Wash the beads once with 400 µL of 50 mM TEAB.
5. Resuspend the beads in 200 µL of 100 mM TEAB and add 4 µg of sequencing-grade trypsin (20 µL of a 0.2 µg/µL solution). Digest overnight (approximately 16 hours) at 37°C with gentle shaking (or in a thermomixer at low speed) (*See* **Note 25**).
6. Centrifuge the samples at 5,000×g for 5 minutes and carefully collect the supernatant containing the tryptic peptides into a new low-protein-binding tube (*see* **Note 26**).
7. Wash the remaining beads twice with 50 µL of 50% ACN/0.1% FA. Pipette up and down to disrupt the bead pellet and extract leftover peptides. Pool these washes with the supernatant from the previous step. Repeat this step once more.
8. Acidify the combined peptide solution to pH ∼3 with 10% formic acid (*see* **Note 27**).
9. Dry the peptides completely in a vacuum concentrator. **STOP POINT**.
10. Desalt the peptides using Pierce Peptide Desalting Spin Columns according to the manufacturer’s protocol.
11. Dry the desalted peptides completely in a vacuum concentrator (*see* **Note 28**). **STOP POINT**.

### 3.9 Tandem Mass Tag (TMT10plex) Labelling

1. Equilibrate the required TMT10plex reagent vials (0.8 mg size) to room temperature before opening (*see* **Notes 29**-**31**). After opening, spin down the reagent in the vial (brief microcentrifuge pulse) to ensure all powder is at the bottom.
2. Resuspend each dried, desalted peptide sample in 100 µL of 100 mM TEAB buffer (*see* **Note 32**).
3. Reconstitute each 0.8 mg TMT reagent vial with 41 µL of anhydrous acetonitrile. Flick or gently vortex the vial to dissolve the reagent; incubate at room temp for 5 min with occasional mixing (*see* **Note 33**).
4. Add the entire 41 µL of the appropriate reconstituted TMT reagent to each corresponding peptide sample. Pipette up and down to mix or gently vortex (*see* **Note 34**).
5. Incubate the labelling reaction for 1 hour at room temperature (*see* **Note 35**).
6. Quench the reaction by adding 8 µL of 5% hydroxylamine solution to each sample. Incubate for 15 minutes at room temperature (*see* **Note 36**).
7. Combine equal amounts from each of the 10 TMT-labelled samples into a single new microcentrifuge tube (*see* **Note 37** and **38**).
8. Dry the pooled sample completely in a vacuum concentrator. **STOP POINT**.
9. Reconstitute the final pooled sample in 100 μL 0.1% formic acid, then desalt to remove TEAB, salts, and residual hydroxylamine using a Pierce Peptide Desalting Spin Column and dry completely in the SpeedVac (*see* **Note 39**). **STOP POINT**.

### 3.10 LC-MS/MS Analysis

*Instrument Setup*: Configure a nanoLC-MS/MS system for data-dependent acquisition (DDA) of TMT-labelled peptides. The following describes typical conditions as used in our analysis, which can be adapted to available instrumentation (*see* **Notes 40**–**42**).

1. Resuspend the final TMT-labelled peptide sample in LC Solvent A (0.1% FA in water).
2. Load the sample onto an Acclaim PepMap 100 C18 trap column (5 μm, 100 µm x 20 mm) and separate on a nanoEase M/Z Peptide BEH C18 analytical column (1.7 μm, 75 µm x 250 mm) using a Thermo Scientific Ultimate 3000 RSLCnano system.
3. Elute peptides using a 150-minute linear gradient from 5% to 35% LC Solvent B (0.1% FA in ACN) at a flow rate of 300 nL/min.
4. Analyse the eluting peptides on an Orbitrap Ascent Mass Spectrometer.
5. Acquire data using the parameters specified in **Table 2**. **STOP POINT**.

**Table 2.** Key LC-MS/MS Acquisition Parameters.

| Parameter | Setting |
| --- | --- |
| MS1 Scan Range | m/z 375–1,500 |
| MS1 Resolution | 120,000 |
| MS1 AGC Target | $4 \times 10^5$ |
| MS1 Max Injection Time | 251 ms |
| MS2 Isolation Window | 0.7 m/z |
| Fragmentation Method | HCD |
| Normalised Collision Energy | 38% |
| MS2 Resolution | 45,000 |
| MS2 AGC Target | 1×10<sup>5</sup> |
| MS2 Max Injection Time | 80 ms |
| MS2 First Mass | m/z 100 |
| Dynamic Exclusion | 45 s |

### 3.11 Data Analysis

1. Process the raw LC-MS/MS data files using MaxQuant software (version 1.6.3.4 or later) with the integrated Andromeda search engine (Cox and Mann, 2008; Cox et al., 2011).
2. Search the spectra against a concatenated database containing both the *L. mexicana* (UniProt KB, 8,559 sequences) and murine proteomes (*see* **Note 43**).
3. Use the search parameters specified in **Table 3**.
4. Process the proteinGroups.txt output file using Perseus software (version 1.6.10.50 or later) (Tyanova et al., 2016). Filter out contaminants, reverse hits, and proteins only identified by site.
5. Normalise the TMT reporter ion intensities by Z-score and transform to a log_2_ scale for quantitative analysis.
6. Perform statistical analysis using a modified *t*-test with permutation-based false discovery rate (FDR, 250 permutations, FDR=0.05) to identify proteins with significant changes in activity between experimental conditions. Apply cutoffs such as, for instance, log_2_ fold-change > 0.5 and –log_10_(p-value) > 0.3 for hits (probe vs. DMSO) (*see* **Note 44**).
7. Visualise the results with a volcano plot: plotting log_2_ enrichment on the x-axis and – log_10_ p-value on the y-axis highlights significant hits. Expect probe targets (active serine hydrolases) to cluster as significantly enriched in the +probe condition.
8. Further bioinformatics (optional): E.g., perform protein family or GO-term enrichment analysis on the hits.
9. Interpretation: Finally, interpret the quantitative results in biological context. Determine which enzymes show increased activity upon infection or which are uniquely detected in the intracellular amastigote stage. For example, in this application INTERACT revealed that 6 parasite serine hydrolases were detectable in infected macrophages (intracellular amastigotes) and that infection induced elevated activity of 5 host hydrolases compared to uninfected macrophages (*vide infra*). These findings point to specific host enzymes possibly co-opted during infection and parasite enzymes critical for intracellular survival. Use appropriate controls to support these interpretations (e.g. uninfected cells with probe, infected with DMSO, etc., as included in the TMT design).

### 3.12 General Notes and Troubleshooting

1. Controls: Always include appropriate controls in ABPP/INTERACT experiments to identify proteins that bind non-specifically (e.g., bead- or tag-associated background). For example, in this study core controls are: • Uninfected macrophages + probe (vs. DMSO) (host targets). • *Leishmania* alone + probe (vs. DMSO) (parasite targets). • Infected + vehicle (DMSO, no probe).
2. In TMT designs, dedicate ≥1 channel to a no-probe (vehicle) control for statistical filtering of non-specific binders. Consider adding beads-only and/or no-click controls when feasible. Without proper controls, abundant proteins (e.g., actin, tubulin) can appear spuriously “enriched”.
3. Safety: Work with live *Leishmania* under BSL-2 practices: wear gloves, lab coat, and eye/face protection; handle all cultures in a Class II biosafety cabinet; and decontaminate waste (e.g., bleach followed by appropriate disposal or autoclaving). Many reagents are hazardous (organic solvents, CuAAC reagents, DMSO-dissolved probes). Use a fume hood for solvent handling and click chemistry. Collect copper-containing waste separately and dispose of it according to institutional regulations (chelating if required). Follow local risk assessments and training requirements.
4. Troubleshooting guidance is summarised in **Table 4**.

## 4. Representative Results

Application of the standardised ABPP protocol with the **FP probe** vs. DMSO successfully identified 14 active serine hydrolases in *L. mexicana* promastigotes, comprising 12 peptidases and 2 lipases. In uninfected murine macrophages, 14 host SHs were also detected as active. Following infection with *L. mexicana*, a marked shift was observed: many host SHs associated with immune defence (including PREP, ABHD12, and Pafah1b3) were no longer detected, suggesting parasite-induced downregulation of these pathways. In parallel, only five host SHs displayed increased activity post-infection.

**Table 3.** Bioinformatic Search Parameters for MaxQuant

| Parameter | Setting |
| --- | --- |
| Type | Reporter ion MS<sup>2</sup> |
| TMT multiplicity | 10-plex TMT |
| Enzyme | Trypsin/P |
| Max. Missed Cleavages | 2 |
| Fixed Modifications | Carbamidomethyl (C) |

| Variable Modifications | Oxidation (M), Acetyl (Protein N-term) |
| --- | --- |
| Peptide FDR | 0.01 |
| Protein FDR | 0.01 |
| Min. Peptide Length | 6 |

**Table 4.** Troubleshooting: common indicators, likely causes, and corrective actions for the INTERACT ABPP/TMT workflow.

| Indicator | Likely cause(s) | Fix / action |
| --- | --- | --- |
| Few or no enriched proteins detected by MS | Probe integrity/dose (FP probe hydrolysis in DMSO; incorrect working concentration). Permeation/incubation suboptimal. CuAAC inefficiency. Matrix effects (detergent/salt before click). | Use fresh aliquots at the correct dose. Confirm entry in both host and parasite by SDS–PAGE; optimise 37°C, up to 1 h, keep DMSO ≤0.1% v/v, verify viability. For CuAAC, run a small pre-enrichment fluorescent gel (e.g., TAMRA-azide). Use fresh CuSO<sub>4</sub> and sodium ascorbate; correct order of addition (ligand present; reduce Cu(II) to Cu(I) *in situ*). Keep lysate conditions as specified; avoid excess detergents/salts pre-click. |
| No/weak fluorescent gel after CuAAC | Degraded click reagents or wrong addition sequence; missing ligand; incompatible buffer conditions. | Replace reagents; ensure ligand present during click; add ascorbate last; verify buffer compatibility; repeat small-scale CuAAC. |
| Weak overall signal, smeared/dirty spectra | SDS/salt carry-over; inadequate cleanup/desalting. | Adhere to stringent wash/desalt steps; if signal depressed or spectra smeared, re-clean samples before MS. |
| Low MS/MS sequence coverage | Suboptimal fragmentation/ acquisition: NCE too low; AGC/injection time too short; isolation window or dynamic exclusion mis-set. | Slightly increase NCE (Normalized Collision Energy) and/or AGC (Automatic Gain Control) target/injection time as duty cycle allows; verify isolation window and dynamic exclusion settings. |
| Low or variable TMT reporter-ion intensities | Incomplete labelling; wrong pH/buffer; moisture/solvent issues; expired/insufficient TMT; short labelling time; high peptide load. | Ensure reagents at RT before opening; reconstitute TMT in anhydrous ACN; peptides in amine-free TEAB, pH 8–8.5; no primary amines (e.g., Tris, glycine). Use sufficient TMT:peptide ratio, mix well, label ≥60 min, quench with hydroxylamine. Store reconstituted TMT briefly at –20°C, avoid moisture; extend time/increase reagent if load was high. |
| Poor TMT quantitation (ratio compression/interference) | Co-isolation and high sample complexity; unequal channel loading; suboptimal isolation/resolution settings. | Fractionate pooled TMT (offline high-pH RP) to reduce complexity; verify equal peptide loading across channels; optimise isolation window, resolution, AGC/injection times. Where available, use SPS-MS<sup>3</sup> to mitigate interference; otherwise tighten isolation with mindful trade-offs and maintain rigorous cleanup. |

Comparison across parasite life stages revealed further changes. While 14 SHs were active in insect-stage promastigotes, only six remained active in the intracellular stage within macrophages during early host–parasite interaction. These enzymes, including prolyl oligopeptidase (POP), oligopeptidase B (OPB), and several lipases, have recognised roles in virulence and lipid metabolism, underscoring their importance for parasite survival and proliferation in the host. Collectively, these findings demonstrate that INTERACT provides a unique snapshot of the host–parasite interplay (**Figure 3**).

**Figure 3.**
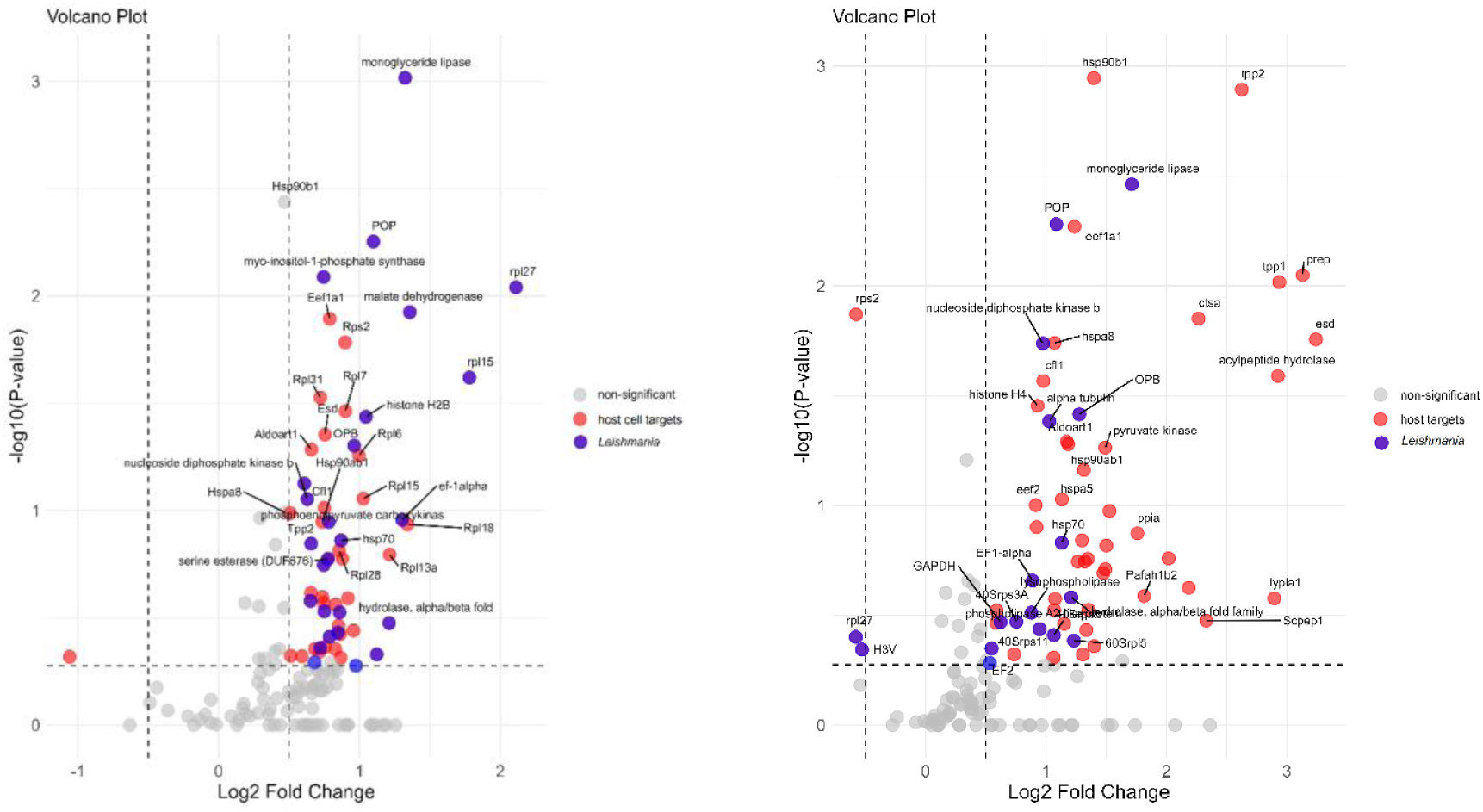
Left: Interactome comparison identifies infection-specific serine hydrolases (SHs) by contrasting infected macrophages (RAW 264.7 cells + *L. mexicana* + **FP probe**) with non-infected controls (RAW 264.7 cells + **FP probe**). Volcano plots show significant enrichment of SHs in the host–parasite interactome at 10 µM **FP probe**; positive targets appear on the right (blue, *L. mexicana*; red, host), while grey points are below significance. **Right:** Probe-oriented comparison in infected macrophages (RAW 264.7 cells + *L. mexicana* + **FP probe**) versus vehicle-treated infected controls (RAW 264.7 cells + *L. mexicana* + DMSO). Data acquired at 10 µM **FP probe** and analysed as described (*see* Section 3.11) reveal significant enrichment of SHs upon probe treatment relative to vehicle. Positive targets appear on the right (blue, *L. mexicana*; red, host); grey points are below threshold.s

The INTERACT workflow is readily transferable to other host–pathogen or host– commensal models with system-specific optimisation. Key variables to adjust include: (1) the activity-based probe to match the enzyme class of interest, (2) culture and infection parameters (e.g., MOI, incubation time, temperature), and (3) capture handle/chemistry when alternative tags are used. TMT multiplexing supports simultaneous comparison of multiple experimental and biological factors (e.g., strains, MOIs, inhibitors, time points), enabling robust, single-experiment contrasts across conditions and positioning INTERACT as a flexible platform for comparative chemoproteomics.

The INTERACT protocol detailed in this chapter represents a pivotal advancement in the study of host-pathogen dynamics, harnessing a powerful chemoproteomic strategy to map enzymatic activity across an interactome under native conditions. By integrating a bespoke, cell-permeable activity-based probe with a quantitative, TMT-based mass spectrometry workflow, this methodology is the first to successfully apply the principles of ABPP for the simultaneous, real-time characterisation of two distinct proteomes during a live infection. This approach overcomes the limitations of traditional lysate-based methods, moving beyond the static analysis of protein abundance to capture the dynamic functional state of enzymes directly *in situ*. As such, this protocol provides a definitive blueprint for researchers seeking to dissect complex intercellular crosstalk, enabling the functional annotation of proteins in their true biological context. It unlocks new and previously inaccessible avenues for understanding infectious disease and is poised to accelerate the discovery of novel therapeutic targets at the host-pathogen interface.

## 5. Notes

1. **FP Probe** Synthesis: This protocol utilizes the cell-permeable Ethyl [5-(prop-2-yn-1-yloxy)pentyl] phosphonofluoridate probe, referred to as **FP Probe**. The synthesis of this probe is detailed in the supplementary information of the primary research article by Isern et al (2025) (named as probe 7) but is beyond the scope of this chapter. The efficacy of this probe stems from a combination of its reactive FP warhead and a linker of sufficient length to ensure its alkyne handle remains solvent-exposed and accessible for the subsequent click chemistry reaction after covalent binding to the target enzyme.
2. Maintain *Leishmania* cultures in exponential growth and avoid overgrowth before differentiation. Parasites that have been passaged too many times in culture can lose virulence so periodic refreshment from frozen stocks is essential. The pH shift to 5.5 in Schneider’s medium is crucial to obtain metacyclic promastigotes. Ensure the medium is properly buffered at pH 5.5 and do not disturb the culture during the 6-day differentiation period. Metacyclic parasites are verified by their agglutination properties or increased infectivity.
3. All INTERACT steps must also be performed on these controls. These controls are essential to distinguish parasite-versus host-derived signals and to validate infection-specific effects.
4. The multiplicity of infection (MOI) can be adjusted depending on experimental goals. An MOI of 10:1 is a good starting point that yields a majority of macrophages infected without excessive free parasites. Lower MOIs (e.g. 5:1) might result in fewer infected cells, whereas very high MOIs can cause macrophage cell stress or death. If macrophage viability is an issue, reduce the MOI or infection time.
5. A 4 h infection allows internalisation of promastigotes (it is short enough to capture early-stage interactions and initial enzyme activation) but limits their transformation into amastigotes. Avoid longer infection times if the goal is to profile initial host– pathogen interactions. If longer infection times are used (e.g. 24 h or more), the parasite will differentiate into amastigotes and replicate, which may change the active proteome. Always consider what stage of infection is being studied and ensure that timing of treatments and lifecycles is consistent across replicates.
6. Thorough washing after infection is critical. Any remaining extracellular *Leishmania* will contribute proteins that are not truly part of the host-internalised interactome. After the third PBS wash, visually inspect the macrophage monolayer under a microscope. There should be little to no motile promastigotes observable in the field with only adherent macrophages visible. If many free parasites are observed, additional washes or gentle pipetting can be undertaken but care is required not to detach macrophages.
7. Match DMSO across all conditions, keeping ≤0.1% v/v if possible. For example, adding 1 µL of probe stock (in DMSO) to 1 mL medium results in 0.1% DMSO, which is generally well tolerated by cells. High DMSO (> 0.5%) can affect cell membrane permeability and enzyme activity, confounding results. If a higher probe concentration is needed, concentrate the probe stock to minimise DMSO volume added.
8. Use a biologically relevant probe incubation temperature. For example, in this case the probe labelling step (Step 3.3.3) is performed at 37°C, which is the optimal (native) temperature for *Leishmania* infection in mammalian host cell. This differs from the 26°C used for *Leishmania* promastigote culture. This is a deliberate choice to maximise parasite enzyme and macrophage activity and ensure robust labelling during this critical step.
9. Protect the labelling incubation from light if using a photo-sensitive probe or if the lab lighting is strong as some warheads can degrade under light. Also, ensure adequate mixing: e.g., in a multi-well plate (or in a flask) which can be achieved by gently tilting or swirling the plate midway through incubation to prevent local depletion of probe near cells.
10. Scraping is used instead of trypsinisation to harvest macrophages because enzymatic detachment (trypsin/EDTA) could potentially digest surface proteins or alter the activity of target enzymes. Additionally, trypsin could remain and later interfere with proteomics. Using cold PBS and a cell scraper preserves protein activity profiles. Work quickly once the plate is out of the incubator; keeping cells cold and proceeding to lysis promptly helps “freeze” the activity state labelled by the probe.
11. The choice of lysis buffer is important. Triton X-100 (1%) is a relatively gentle non-ionic detergent that preserves protein complexes and is compatible with downstream steps. Also, including glycerol (5%) helps stabilise proteins and reduce aggregation during lysis.
12. Ensure complete removal of insoluble debris after centrifugation. The pellet may contain unbroken nuclei, cell walls, or unsolubilized membranes. If the pellet is loose and / or sample loss is suspected, re-lyse the pellet in a small volume of lysis buffer and combine with the supernatant. The clarity of the lysate is crucial for efficient downstream click chemistry and binding; any cloudiness might indicate remaining lipids or particulate matter, which can clog spin columns or bind non-specifically to beads. This solution can be stored at −80°C for several months.
13. Click reagent addition: For the click reaction (Step 3.6), it is critical to add the reagents in the correct order to ensure efficient catalysis. The protein lysate, Biotin- N_3_, CuSO_4_, and the copper-chelating ligand (TBTA) should be mixed first. The reaction is initiated by the addition of the reducing agent (sodium ascorbate), which generates the catalytic Cu(I) species *in situ*.
14. The reaction is aqueous but contains some DMSO from stocks; ensure this remains ≤ 5% total
15. During this time, the alkyne-labelled proteins react with the azide-biotin, appending a biotin tag to each probe-bound enzyme. Periodically invert or gently vortex the tubes to keep the solution homogeneous. Optional: Quench the reaction with EDTA at 5 mM (final concentration).
16. If sample volume is large, it is acceptable to split the reaction mixture into multiple tubes for precipitation. For better results, overnight incubation is preferred.
17. Do not over-dry to the point of a hard pellet; a brief dry is sufficient as any remaining moisture will be addressed in re-solubilisation.
18. This yields ∼1 mL of diluted lysate per sample. The SDS helps solubilise proteins and the dilution ensures the final SDS concentration is low enough for binding to beads. SDS is necessary to solubilise the proteins, but too much SDS can be problematic later. By immediately diluting to 0.1% SDS. This is sufficient to keep proteins soluble and is tolerable for NeutrAvidin binding (NB NeutrAvidin is quite stable and can handle some detergent). However, more than 0.2% SDS can start to elute proteins non-specifically from the beads or denature the binding too much. The extensive washing later (with SDS, urea, etc.) further ensures that only biotin-tagged proteins stay attached.
19. This allows biotin-tagged proteins to bind to the NeutrAvidin (or streptavidin) on the beads. Ensure continuous gentle mixing so beads do not settle.
20. Affinity enrichment washes: The sequential, stringent wash steps (Step 3.7.7) are crucial for removing non-specifically bound proteins and residual reagents, thereby reducing background and improving the signal-to-noise ratio in the final MS analysis. The use of both a strong detergent (SDS) and a chaotropic agent (urea) ensures high-purity enrichment of biotinylated proteins. Perform each wash by adding buffer, gently inverting the tube several times (or flicking), then pelleting beads (1,500 × g, 2 min) and removing supernatant. Combine the quickness of washes with thorough mixing to maximise removal of contaminants.
21. To remove any remaining hydrophobic or weakly bound proteins.
22. To disrupt strong non-specific interactions and denature proteins.
23. To remove urea and bring beads back to a mild buffer.
24. To exchange into a digestion-compatible buffer.
25. This is the first peptide fraction.
26. On-bead digestion workflow: The on-bead digestion method (Step 3.8) is a critical component of the workflow for generating a clean peptide sample. This approach ensures that only peptides derived from the captured, probe-labelled proteins are released into the solution for subsequent analysis. Attempting to elute the intact proteins from the beads before digestion is not recommended, as this can lead to significant sample loss, increased background from non-covalently bound contaminants, and lower overall sensitivity. On-bead digestion often yields slightly fewer peptides than in-solution digestion of the same protein, because diffusion to the enzyme is somewhat hindered. To maximise yield it is acceptable to increase the trypsin amount (e.g. in this case 4 µg was used for <100 µg protein on beads) and digestion time. Mixing is also important. If using a thermomixer, ∼800 rpm at 37°C is ideal. If not, gently tapping or flicking the tube occasionally during incubation will suffice (or place on a rotating mixer in a 37°C room). After overnight digestion, the liquid should be clear of the viscous bead slurry obtained when you spin down the beads.
27. The acidification stops trypsin activity and improves binding to C18 during desalting.
28. The result is a dried peptide sample enriched for probe-labelled proteins (store at - 80°C, if not used immediately).
29. TMT reagent handling: The TMT reagents are highly moisture-sensitive. To avoid hydrolysis, which deactivates the reagent and leads to poor labelling efficiency, the reagent vials must be fully equilibrated to room temperature before opening. Use anhydrous ACN for reconstitution. Unused reconstituted reagent can be stored at - 20°C for up to one week but must be warmed to room temperature before opening for subsequent use.
30. When transferring the dissolved TMT reagent, be cautious of glass vial surfaces. TMT reagents can stick to vial walls if not fully dissolved. If possible, after adding ACN and dissolving, use the same vial as the reaction container by adding the peptide solution into it. This ensures quantitative transfer of the tagging reagent. Also, perform a quick check that the pH of the peptide solution is around 8 by spotting a tiny aliquot on pH paper (too acidic or too basic can reduce labelling efficiency). TEAB at 100 mM usually suffices to maintain pH in the optimal range.
31. Amine-Free Buffers for TMT Labelling: The TMT labelling reaction targets primary amines on the N-termini of peptides and the side chains of lysine residues. It is therefore essential that all buffers used during and after the tryptic digest (Steps 3.8 and 3.9) are free of primary amines (e.g., Tris, glycine), which would compete with the peptides and quench the reaction. The use of TEAB buffer is required for this reason.
32. It is recommended to have all samples at the same peptide concentration for equal labelling efficiency. Optionally, verify peptide concentration with a quantitative peptide assay (avoid assays with primary amines).
33. Check that the solution is clear (without undissolved particles).
34. Alternatively, transfer the peptide solution directly into the TMT vial to ensure complete transfer of reagent. Label each sample with a different TMT channel. Record which sample gets which reagent.
35. The reaction covalently attaches the TMT tag to peptide N-termini and lysine side- chains. Ensure the pH remains ∼8–8.5 during labelling (TEAB buffer should maintain this; if the solution was acidic, the reaction efficiency drops). If precipitation is observed (rare), dilute with more TEAB buffer.
36. This will neutralise any unreacted TMT reagents, preventing them from labelling peptides later when samples are combined.
37. Ideally, base this on peptide amount rather than volume; you may measure peptide concentration post-labelling by a colorimetric assay that is compatible with TMT (the BCA peptide assay cannot be used after TMT labelling, but an amino acid analysis or a calibrated LC-MS signal could be used). In absence of a precise measurement, assume equal recovery and combine equal volumes. Gently vortex the combined sample.
38. It is critical to combine equal peptide amounts from each TMT channel. Any imbalance will directly affect the accuracy of relative quantitation. If one channel (e.g. a control) has much lower peptide content, it may result in missing values or lower signal for that channel, which could be misinterpreted as a real difference. Starting with equal quantities of protein and, assuming equal digestion efficiency, combining equal volumes is normally an acceptable approximation. For more accuracy, analyse a small aliquot of each labelled sample on LC-MS to quantify a common peptide and normalise, but this is often not necessary with careful workflow.
39. Store dry TMT-labelled peptide mixture at –80°C if not proceeding immediately to LC-MS. At this stage, the sample represents the combined tryptic peptides of all probe-labelled proteins from all conditions, differentially tagged for relative quantitation.
40. To ensure consistent peptide fragmentation and reduce redundant sampling, standard data-dependent acquisition (DDA) filters were applied. Monoisotopic precursor selection was enabled; precursors with unassigned charge or charges outside the typical peptide range were excluded from fragmentation; and an instrument-appropriate minimum intensity threshold was used for MS/MS triggering. To limit repeat sequencing, dynamic exclusion was used (single count, ∼30–60 s duration) with isotope exclusion and a narrow mass-tolerance window (±5–10 ppm). Final values should be tuned to instrument model, chromatographic peak width, and sample complexity.
41. LC-MS/MS fragmentation: Higher-energy collisional dissociation (HCD) is required for the fragmentation of TMT-labelled peptides to generate the low-mass reporter ions used for quantification. ETD alone does not efficiently generate low-mass TMT reporter ions; use HCD (or HCD-MS^3^ where available). Ensure the mass spectrometer is configured for HCD fragmentation and that the MS^2^ resolution is set to at least 45,000 to resolve the isobaric reporter ions effectively.
42. Achieving high resolution in MS^2^ is key for TMT. Here, 45k resolution was used which cleanly separates the 1 Da apart reporter ions. Lower resolution (e.g. 15k) can cause overlap of reporter ion peaks, leading to inaccuracies. If using an older mass spectrometer that is not capable of high-resolution MS^2^, consider using MS^3^-based methods (like SPS MS^3^ on Orbitrap Fusion) to circumvent co-isolation interference. Optimal normalised collision energy (NCE) can vary (e.g. 35%–40%). Whilst the recommended NCE (∼38%) was optimal in this example tuning may be required. If reporter ions are weak then NCE is too low whilst if peptide sequence ions are fragmented to noise then NCE settings are too high. Tuning with a TMT-labelled standard (Thermo provides a TMT tuning mix) can help.
43. Using a combined database for host and pathogen is important to avoid misidentification. If a peptide sequence is shared between a host and parasite protein, database search algorithms might randomly assign it to one or the other. MaxQuant’s “matching between runs” can propagate identifications across LC-MS runs, but in a single TMT multiplex run this isn’t applicable. To increase confidence, undertake separate searches against each organism and compare. The simplest method is a combined FASTA with organism prefixes and parsing using those prefixes. In this analysis, all *Leishmania* proteins had accessions like “E9xxxx” and gene codes “LmxM.”, whereas the mouse proteins have gene names or UniProt IDs that are distinct making it simple to separate them post-analysis.
44. The statistical cutoffs should be adjusted based on the number of replicates and data variability. Biological duplicates for each condition, which, combined with moderated *t*-test, gives confidence in identifying hits. Although not ideal, if only one replicate per condition is possible consider using fold-change ranking rather than strict p-values. If missing values occur in the TMT analysis, for example when a protein is only identified in probe-treated samples and not in controls (or vice versa), impute a low intensity for the missing channel or treat that protein qualitatively as “only in probe”. This can result in some targets only appearing on one side of the volcano plots. Perseus software can perform imputation (e.g. drawing from a low-intensity distribution) to allow log_2_ fold-change calculation.

## Acknowledgements

The authors thank the UKRI-Global Challenges Research Fund. A Global Network for Neglected Tropical Diseases MR/P027989/1 (to P.G.S. and P.W.D.); The Royal Society (The Royal Society International Collaboration Awards for Research Professors 2016: IC160044 (to P.G.S.); MRC IAA 2021 Durham University MR/X502947/1 (to P.W.D.); COFUND (European Union and Durham University) (Junior Research Fellowships (JRF) to E.O.J.P.) for financial support.

